# The Survival of β-lactoglobulin Peptides in the Archaeological Record: Vulnerability vs. Sequence Variation

**DOI:** 10.1101/2024.09.13.612646

**Authors:** Beatriz Fonseca, Colin L. Freeman, Matthew J. Collins

## Abstract

It is a strange observation, given the cultural co-evolution of dairying, that milk proteins are more commonly reported than any other food proteins in the archaeological record. The whey protein β-lactoglobulin and in particular its eleven amino acid long peptide T_125_PEVDXEALEK_135_ seems to be preferentially preserved in both ceramic vessels and teeth (dental calculus). An amino acid substitution in the middle of the chain is valuable to track livestock management because it permits differentiation between key animal species used in dairying. The persistence of this peptide is, however, unusual as its acidic nature makes it more vulnerable to hydrolysis. Moreover, selection for the ability to digest raw milk - more specifically, the continued production of the milk sugar enzyme lactase beyond the age of normal weaning - did not begin until the early Bronze Age. It is therefore unclear why it is a milk peptide, in particular a peptide associated with the lactose-rich whey fraction, that is one of the most commonly recovered dietary peptides. The unexpected preservation of T_125_PEVDXEALEK_135_ thus presents a good case study to uncover patterns of protein survival in the archaeological record. We have previously explored the dynamics of the bovine variation of the peptide (X=Asp_130_) and its likelihood to undergo hydrolysis in solution. In this study, we turn our attention to the ovine (X=Asn_130_) and the caprine (X=Lys_130_) variations of the β-lactoglobulin peptide to determine how the mutation in the amino acid in position 6 affects peptide conformations and vulnerability in bulk water. To do this, we use Molecular Dynamics as implemented in GROMACS 2020, with the Amber14SB forcefield and the SPC/E water model. We first perform extensive conformational analysis of both peptides in solution to determine stable structures. Then, using analogous methodology to that developed in our earlier study of the bovine peptide, we identify geometric arrangements between water and peptide that may be more prone to hydrolysis.

## Introduction

Following the first reports of milk proteins in food crusts from the iconic early Neolithic site of Çatalhöyük^1^, proteomics has been combined with lipid residue analysis to explore evidence of dairy exploitation, not least because the ability to consume raw milk (specifically the milk sugar lactose) is the most strongly selected trait in modern humans. Milk proteins are amongst the most commonly reported protein residues from dental calculus and limescale deposits in ceramic pots^1–6^. The whey protein β-lactoglobulin (BLG), which partitions with milk sugars when milk is curdled, seems to be preferentially preserved^4,6^. Of all β-lactoglobulin peptides, T_125_PEVDXEALEK_135_ is one of the most commonly recovered and has been reported in several studies on dairying practices^3,4,6,7^. This peptide presents a particular advantage for the study of herding and dairying in past societies, as the substitution in the middle of the chain allows for differentiation between common animals used for this purpose, i.e. cow (Asp_130_), sheep (Asn_130_) and goat (Lys_130_) ^2–4,7^. From this point forward, these animals will be referred to as bovine, ovine and caprine, respectively (see Table S1 for details).

The survival of the whey peptide T_125_PEVDXEALEK_135_ is unexpected for two reasons. Firstly, because early populations were unable to digest lactose^5,8,9^, curdling or fermentation techniques had to be employed to remove the lactose from the whey fraction of the milk. Secondly, the peptide T_125_PEVDXEALEK_135_ is rich in acidic residues and comes from a loop region of the protein, which is naturally more vulnerable to degradation^10–13^. In the study of ostrich eggshell proteins, similar acidic peptides have been found to break down in solution and their survival has been linked instead to the presence of a mineral surface^14,15^. This same BLG sequence has also been found to preferentially bind to metal surfaces in equipment used in the dairy industry^16,17^. The mineral matrix may then play an important role in the persistence of T_125_PEVDXEALEK_135_. This concept of the role of mineral surfaces in enhancing the survival of sequences widely reported^14,15,18^, but outside the study of ostrich eggshell, there is little direct evidence for this supposition.

To understand the unlikely survival of T_125_PEVDXEALEK_135_, it is first necessary to determine its dynamics and vulnerability to hydrolysis in solution. The hydrolysis of a peptide bond has been largely studied in the literature^19–34^ and a few reaction mechanisms have been proposed^10,20,21,25,32^. In our previous work, we have extensively discussed these mechanisms and used published literature values of reactant structures to identify vulnerable peptide bonds along the bovine T_125_PEVD**D**EALEK_135_ peptide chain^19^. Although chemical reactions are better described by quantum mechanical (QM) methods, the size of the system makes it prohibitive to use such techniques. We have therefore successfully developed a methodology to explore the susceptibility of a peptide bond to undergo hydrolysis using classical molecular dynamics (MD)^19^.

Now, in order to have a more detailed picture on the unusual preservation of T_125_PEVDXEALEK_135_, we seek to understand how the substitution in position 6 of the peptide affects its dynamics. To do this, we perform an analogous study to that of the bovine T_125_PEVD**D**EALEK_135_ in solution on the ovine (T_125_PEVD**N**EALEK_135_) and caprine (T_125_PEVD**K**EALEK_135_) variations of the β-lactoglobulin peptide.

## Methods

### Simulation details

The initial structure for the ovine T_125_PEVD**N**EALEK_135_ and caprine T_125_PEVD**K**EALEK_135_ peptides were taken from the corresponding full β-lactoglobulin protein (PDB ids: 4CK4^35^ and 4OMX^36^, respectively). All simulations were performed using the GROMACS 2020.3^37–39^ Molecular Dynamics software.

In our previous analogous work on the bovine T_125_PEVD**D**EALEK_135_ peptide^19^, the system was modelled at pH 7, with the amine and carboxylic groups in their charged form (NH_3_^+^ and CO_2_^-^) and the N- and C-terminal uncapped. For comparison purposes, the systems with the caprine and ovine peptides were run in the same conditions and with the same Molecular Dynamics parameters, described as follows. Equilibration of the system was first performed for 1 ns in the NVT ensemble with a reference temperature of 300 K, followed by a 1 ns run at the NPT ensemble with a reference pressure of 1 bar. The Parrinello-Rahman barostat, with a time coupling constant of 0.2 ps, was used to control the pressure. After equilibration, MD production runs were performed in the NVT ensemble at 300 K, with a time step of 0.5 fs. For temperature control, a Nosé-Hoover thermostat, with a time coupling constant of 0.1 ps, was used. Long range electrostatic interactions were calculated with the Particle Mesh Ewald (PME) method with a cut-off of 1.0 nm, Fourier grid spacing of 0.12 nm and a 4^th^ order spline. A cut-off distance of 1.0 nm was set for Lennard-Jones interactions. The peptide was described by the Amber14SB^40^ forcefield and solvated with 5676 water molecules described by the SPC/E^41,42^ model, due to its good reproducibility of water dynamics^41^. The system consisted of a cubic box with periodic boundary conditions applied in all directions. Na^+^ ions were added accordingly to neutralise the net charge of the system (- 3*e* for the ovine peptide and -2*e* for the caprine peptide).

To enhance sampling of the conformation space of the peptides, metadynamics^43–45^ (metaD) was performed using the radius of gyration (R_g_) as a collective variable, given the flexible nature of the peptides being studied. Gaussian hills with a width of 0.35 and height of 1 kJ/mol were added at an interval of 0.5 ps. Runs of 30 ns in length were performed with the PLUMED^46,47^ v2.6 plug-in, coupled with GROMACS.

### Exploring conformational space

To have a grasp on the impact of the amino acid in position 6 on the number of conformations a given peptide may be able to adopt, the flexibility of each residue in the amino acid chain was investigated by calculating the average root mean square fluctuation (RMSF) along a 50 ns MD trajectory, before conformational analysis. The conformational space of both peptides was then explored using a combination of metadynamics and standard molecular dynamics simulations, following the methodology published in Fonseca et al^19^. To ensure good statistics to explore water dynamics and potential sites for hydrolysis, a given peptide conformation was considered stable when it was maintained for at least 5 ns with a maximum variation of 0.2 nm in the root mean square deviation (RMSD) of the backbone atoms (C Cα N O) during a MD simulation. Its contribution to the total average potential energy ⟨*E*⟩ was then calculated using a Boltzmann probability distribution. When there were no significant changes to ⟨*E*⟩, the conformational space was considered sufficiently sampled. The conformations with major contributions to ⟨*E*⟩ were selected for further analysis.

### Water dynamics and potential sites for hydrolysis

To investigate water dynamics around the peptide bonds, the autocorrelation functions for hydrogen bonds between each peptide bond and water were computed following the method published by Luzar & Chandler^48,49^. The same method was applied to calculate the autocorrelation functions for hydrogen bonds between only the carbonyl oxygen of the peptide backbone and the hydrogen of water H_w_, given that the carbonyl is a primary site of attack for hydrolysis. Results were then combined with the occurrence of geometric parameters for reactant structures published in the literature^20,21,25^, using the same procedure employed for the corresponding bovine β-lactoglobulin peptide in solution in our previous work^19^.

For comparison purposes, relevant data for the bovine T_125_PEVD**D**EALEK_135_ peptide was taken from Fonseca et al^19^, in order to have a more complete picture of the influence of the amino acid in position 6 on the dynamics of the β-lactoglobulin peptides in solution and their likelihood to undergo hydrolysis.

All analyses were performed using GROMACS standard analytical tools or scripts written in Python 3 by the authors.

## Results and Discussion

### Peptide flexibility and conformational analysis

The T_125_PEVDXEALEK_135_ peptide is a very flexible amino acid chain, given the acidic nature of the sequence combined with its small size and overall charge (-4*e* for the bovine peptide, -3*e* for the ovine peptide and -2*e* for the caprine peptide at neutral pH). Figure 1 showcases the average root mean square fluctuation (RMSF) of each residue, for each of the peptide variations, along a 50 ns molecular dynamics simulation. Overall, flexibility tends to increase towards the N- and the C-terminals. In the bovine peptide, for example, residue flexibility is higher in each of the terminals than in the rest of the chain, although Pro_2_ and Glu_3_, towards the N-terminal, have the highest fluctuation overall when compared to these same amino acids in the other peptide variants. For the ovine and the caprine peptides, there are differences in flexibility in either side of the mutation amino acid in position 6. The ovine peptide has the lowest fluctuation overall, which suggests a lower degree of freedom when compared to the other two. When comparing both sides of the ovine sequence, the charged residues from the middle of the chain towards the C-terminal display increased mobility. This trait is particularly marked in the last two amino acids: the glutamate in position 10 and the C-terminal lysine. The opposite trend is observed in the caprine peptide, where the flexibility is slightly increased from the glutamate in position 7 towards the N-terminal instead. The difference in charge distribution in each of the peptides, caused by the substitution in the amino acid in position 6, gives rise to these different trends. As anticipated, the mutation therefore has an impact on the behaviour of the peptide and potentially on the different types of conformations it can adopt.

**Figure 1.**
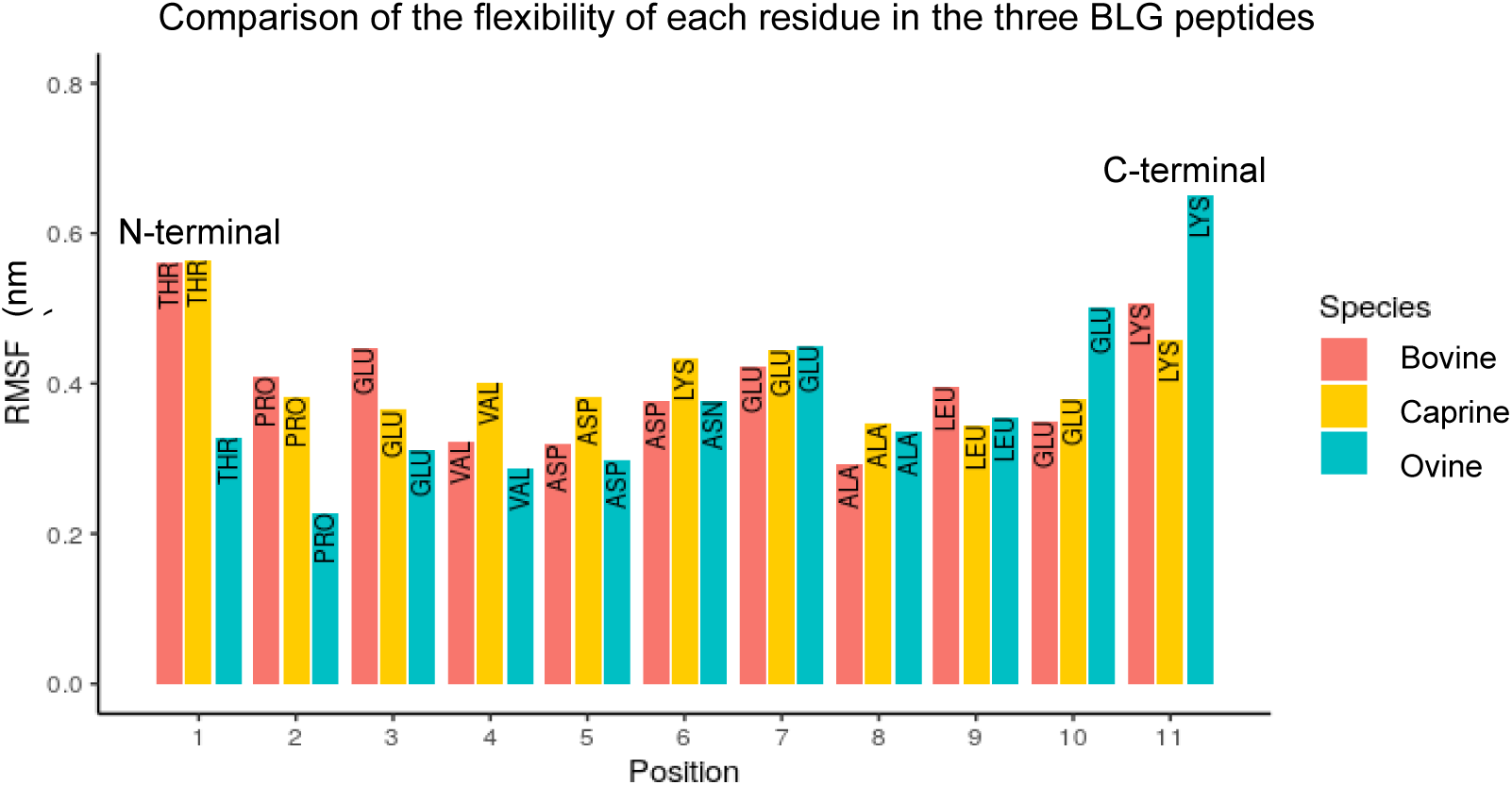
Root mean square fluctuation of each residue in each BLG peptide variation.

Metadynamics simulations revealed that the free energy surface (FES) along the radius of gyration coordinate (Figure 2) has two distinct minima for both the caprine and the ovine peptide, which refer to a folded and an unfolded conformation (Table 1). The bovine peptide, however, has very broad minima, for which the energy barrier is almost non-existent. For all three peptides, the first minimum refers to a folded conformation, whereas the second minimum refers to an unfolded conformation. The first minimum of the ovine peptide, in particular, has a lower energy than the unfolded conformation. Additionally, the barrier to cross to an unfolded state is higher than to cross from an unfolded state back to a folded one. Folded conformations are then favoured for T_125_PEVD**N**EALEK_135_. In the case of the caprine peptide T_125_PEVD**K**EALEK_135,_ the second minimum has a slightly deeper energy well than the first minimum with barriers of similar height in both directions, with a slight advantage towards an unfolded conformation.

**Figure 2.**
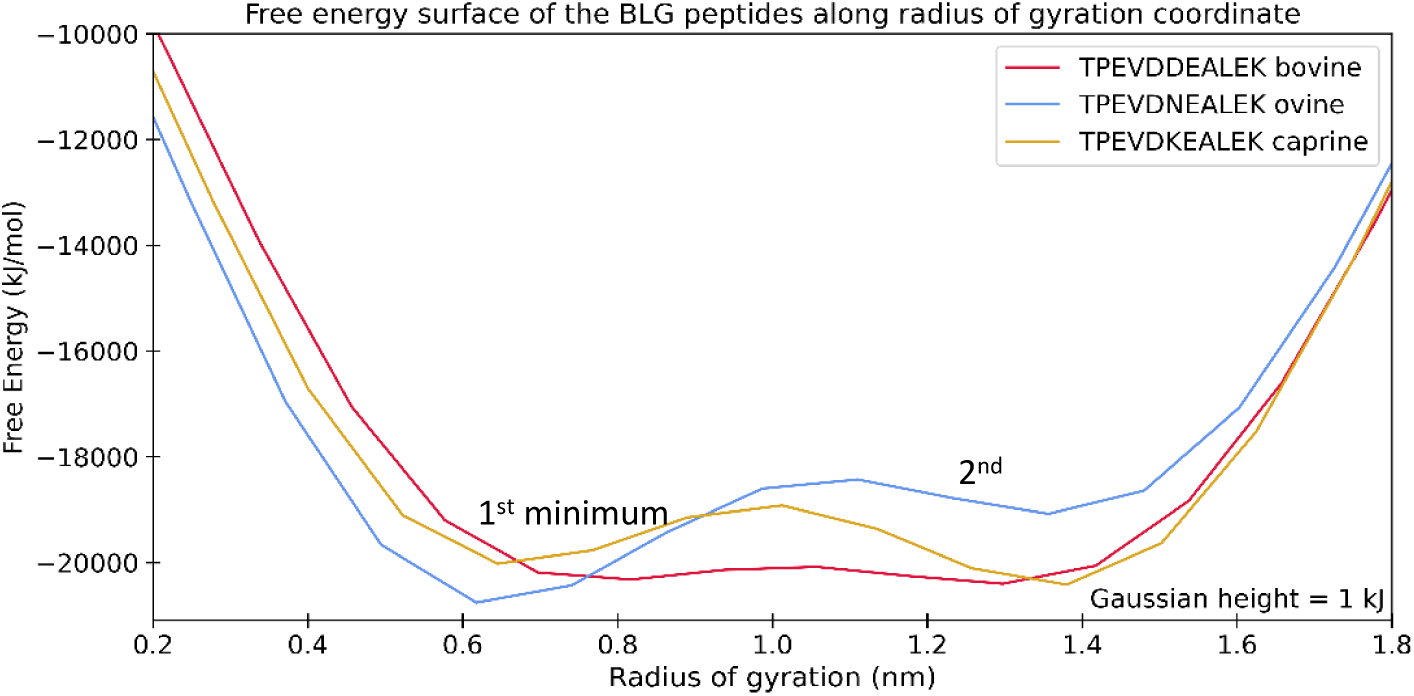
Free energy surface of the bovine β-lactoglobulin T_125_PEVD**D**EALEK_135_ (red), the ovine β-lactoglobulin T_125_PEVD**N**EALEK_135_ (blue) and the caprine β-lactoglobulin T_125_PEVD**K**EALEK_135_ (yellow) peptides obtained by performing metadynamics along the radius of gyration coordinate. The curve for the bovine β-lactoglobulin T_125_PEVD**D**EALEK_135_ was obtained from our previous work (see Fonseca et al.^18^).

**Table 1.**
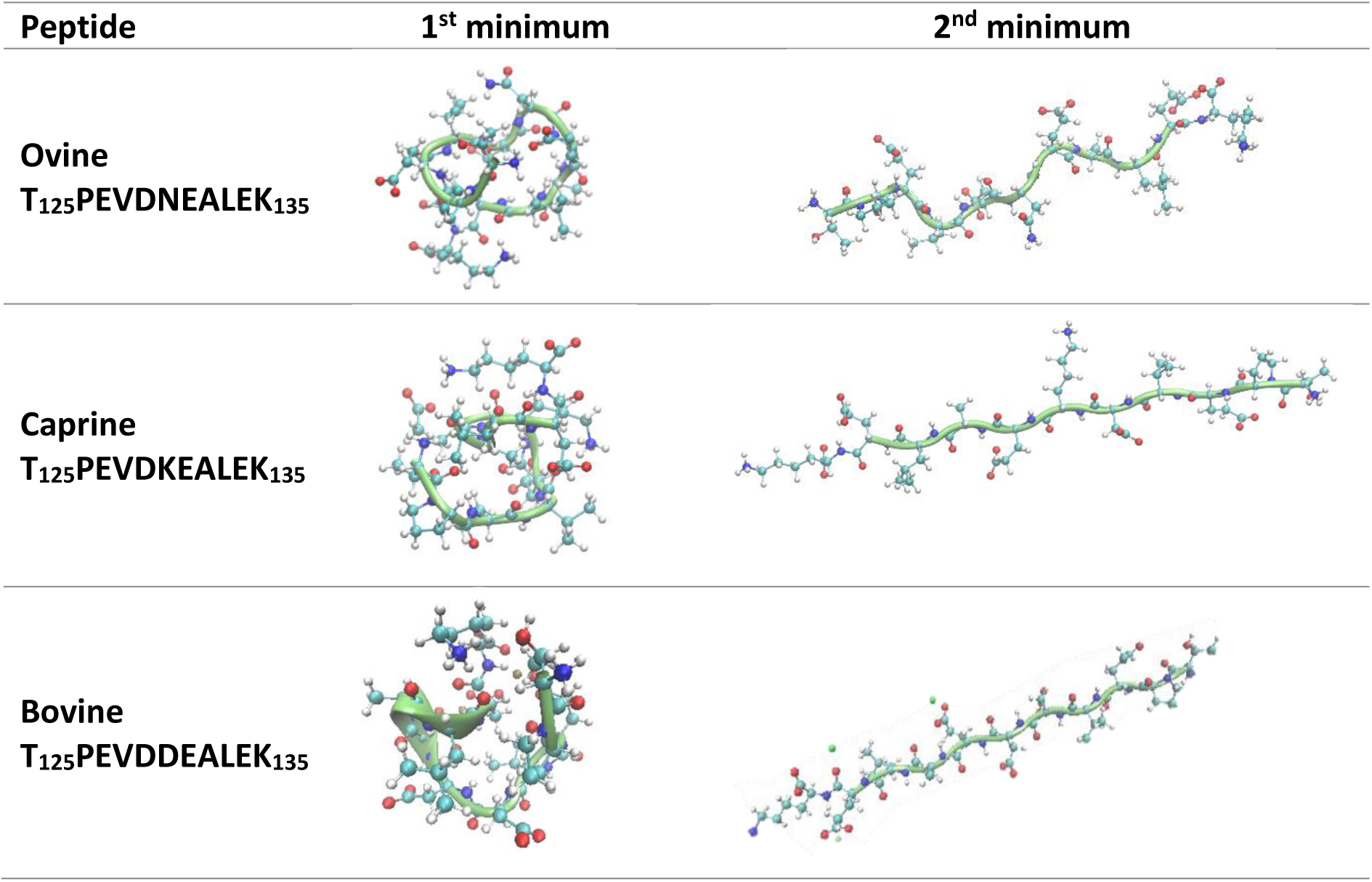
Structures corresponding to the energy minima of the free energy surface along the radius of gyration coordinate, as determined by metadynamics, for the three BLG peptides. For comparison purposes, structures for the bovine peptide were taken from Fonseca et al.^19^

| Peptide | 1<sup>st</sup> minimum | 2<sup>nd</sup> minimum |
| --- | --- | --- |
| Ovine<br>T<sub>125</sub>PEVDNEALEK<sub>135</sub> |  |  |
| Caprine<br>T<sub>125</sub>PEVDKEALEK<sub>135</sub> |  |  |
| Bovine<br>T<sub>125</sub>PEVDDEALEK<sub>135</sub> |  |  |

Conformational analysis unveiled three dominant structures for the ovine peptide and four dominant structures for the caprine peptide (Figure 3). The lower number of stable structures for the ovine variant is consistent with the metadynamics results (these showed a higher energy barrier between states) and with the reduced flexibility of its residues. By contrast, the bovine peptide can easily switch between a folded and an unfolded state, and our previous work has found five stable structures for this variant^19^. It is noteworthy that, for the ovine peptide, the dominant folded structure takes the form of a helix (Structure 1s) instead of the coil type formation seen in metadynamics. This difference is probably because the radius of gyration was chosen as a collective variable (CV). A combination of the torsion angles phi *φ* and psi *ψ* may be a more adequate CV to identify the helix conformation on the free energy surface. Metadynamics was only used as a secondary tool to aid in the exploration of the conformational space of the BLG peptides and not intended to define the whole energy landscape. The determination of the free energy surface along other geometric coordinates was therefore not within the scope of this work. Nonetheless, the recurrence of a helix in the unbiased MD simulations for both the ovine and the caprine peptides - and the fact that it is a type of folded conformation - suggests that this is a stable geometric arrangement for these sequences.

**Figure 3.**
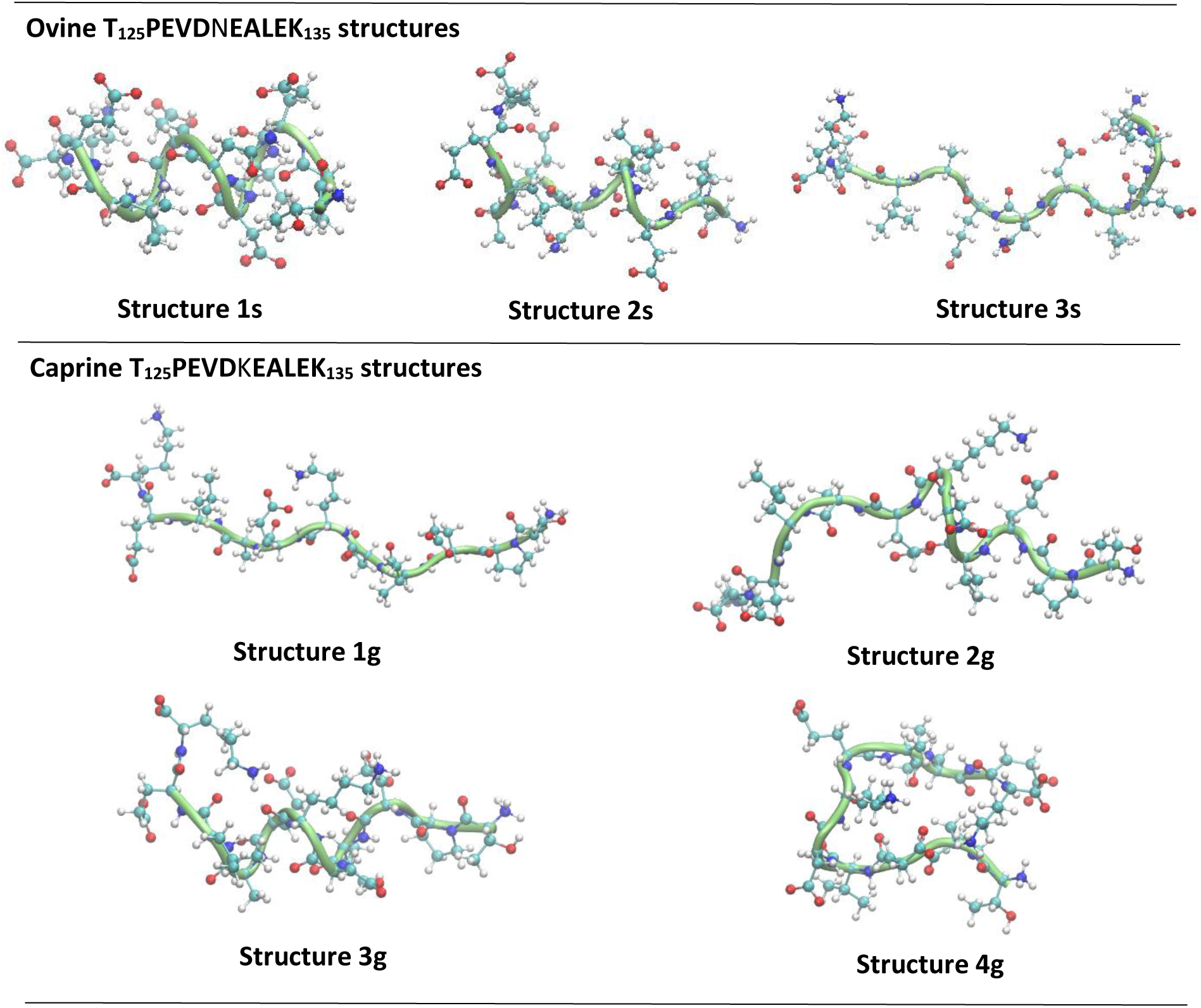
Stable conformations of the TPEVD**N**EALEK ovine (top) and TPEVD**K**EALEK caprine (bottom) β-lactoglobulin peptides in solution, found in replica and metadynamics simulations.

For the ovine peptide T_125_PEVD**N**EALEK_135_, the three stable structures consist of a helix (Structure 1s), a partial helix (Structure 2s) - where part of the chain forms a helix type turn - and a stretched conformation with a small coil towards the N-terminal of the sequence (Structure 3s). In the case of the caprine peptide T_125_PEVD**K**EALEK_135_, the four stable structures consist of a stretched conformation (Structure 1g), a partial helix (Structure 2g), a helix (Structure 3g), and a coil (Structure 4g) (Figure 3). It is interesting to note that, unlike several of the structures found for the bovine peptide^19^, none of the structures for the ovine and caprine variants rely on structural Na^+^ to maintain their conformation. They rely on intramolecular hydrogen bonds instead. This effect is a consequence of the different charge distribution along the peptide sequence caused by the substitution in the amino acid in position 6. For the bovine peptide, T_125_PEVD**D**EALEK_135_, the mutant amino acid is an aspartate (D) and the middle of the sequence is then composed of three acidic residues: DDE. As a consequence, this region in the sequence has a high density of negative charge, which add to repulsive forces in the chain that need to be stabilised either by interaction with the side chain of the N-terminal lysine or by Na^+^ ions^19^. In the case of the ovine and the caprine peptides, the mutant amino acids are asparagine (N) and lysine (K), respectively. Whilst the asparagine has an amide in the side chain, the lysine has a positively charged amine (at neutral pH), both of which help stabilise the negatively charged aspartates and glutamates present in the sequence by helping to minimise the repulsive forces between the carboxyls. This is evidenced by the higher number of intramolecular hydrogen bonds the substituted residues form with the rest of the chain in structures that show some degree of folding (see Table 2 for the ovine peptide and Table 3 for the caprine peptide). This effect is observed even for Structure 1g, the extended conformation for T_125_PEVD**K**EALEK_135_. In this case, the Lys_6_ side chain hydrogen bonds with the glutamate in position 7, which helps to keep the conformation unfolded.

**Table 2.** Average number of intramolecular hydrogen bonds per residue in the ovine BLG peptide T_125_PEVDNEALEK_135_.

| RESIDUE | STRUCTURE 1s<br>(helix) | STRUCTURE 2s<br>(partial helix) | STRUCTURE 3s<br>(stretched) |
| --- | --- | --- | --- |
| THREONINE 1 | 0.15 ± 0.01 | 0.16 ± 0.00 | 0.95 ± 0.02 |
| PROLINE 2 | 1.07 ± 0.01 | 0.85 ± 0.02 | 0.00 ± 0.00 |
| GLUTAMATE 3 | 0.94 ± 0.01 | 0.59 ± 0.02 | 0.20 ± 0.02 |
| VALINE 4 | 0.36 ± 0.01 | 0.06 ± 0.01 | 0.34 ± 0.02 |
| ASPARTATE 5 | 0.88 ± 0.01 | 0.83 ± 0.02 | 0.62 ± 0.02 |
| ASPARAGINE 6 | 1.73 ± 0.01 | 2.01 ± 0.03 | 0.27 ± 0.02 |
| GLUTAMATE 7 | 1.04 ± 0.01 | 0.33 ± 0.02 | 0.24 ± 0.02 |
| ALANINE 8 | 0.55 ± 0.01 | 0.77 ± 0.01 | 0.11 ± 0.01 |
| LEUCINE 9 | 0.62 ± 0.01 | 0.59 ± 0.02 | 0.07 ± 0.01 |
| GLUTAMATE 10 | 0.25 ± 0.01 | 0.10 ± 0.01 | 0.03 ± 0.01 |
| LYSINE 11 | 0.53 ± 0.01 | 0.64 ± 0.03 | 0.05 ± 0.01 |
| TOTAL PER<br>STRUCTURE | 8.12 ± 0.03 | 6.93 ± 0.06 | 2.88 ± 0.05 |

**Table 3.** Average number of intramolecular hydrogen bonds per residue in the caprine BLG peptide T_125_PEVDKEALEK_135_.

| RESIDUE | STRUCTURE 1g<br>(stretched) | STRUCTURE 2g<br>(partial helix) | STRUCTURE 3g<br>(helix) | STRUCTURE 4g<br>(loop) |
| --- | --- | --- | --- | --- |
| THREONINE 1 | 0.01 ± 0.00 | 0.11 ± 0.01 | 0.02 ± 0.00 | 0.08 ± 0.01 |
| PROLINE 2 | 0.00 ± 0.00 | 0.69 ± 0.02 | 0.68 ± 0.01 | 0.08 ± 0.01 |
| GLUTAMATE 3 | 0.55 ± 0.02 | 1.26 ± 0.03 | 0.94 ± 0.01 | 0.95 ± 0.02 |
| VALINE 4 | 0.51 ± 0.02 | 0.46 ± 0.02 | 0.55 ± 0.01 | 0.22 ± 0.01 |
| ASPARTATE 5 | 0.15 ± 0.01 | 0.59 ± 0.02 | 0.73 ± 0.01 | 0.39 ± 0.01 |
| LYSINE 6 | 0.42 ± 0.02 | 1.42 ± 0.03 | 1.46 ± 0.01 | 1.03 ± 0.02 |
| GLUTAMATE 7 | 0.19 ± 0.01 | 0.84 ± 0.02 | 1.13 ± 0.01 | 0.04 ± 0.00 |
| ALANINE 8 | 0.01 ± 0.00 | 0.24 ± 0.02 | 0.66 ± 0.01 | 0.07 ± 0.01 |
| LEUCINE 9 | 0.11 ± 0.01 | 0.09 ± 0.01 | 0.70 ± 0.01 | 0.05 ± 0.00 |
| GLUTAMATE 10 | 0.12 ± 0.01 | 0.02 ± 0.01 | 0.32 ± 0.01 | 0.39 ± 0.01 |
| LYSINE 11 | 0.18 ± 0.01 | 0.04 ± 0.01 | 0.61 ± 0.01 | 0.50 ± 0.02 |
| TOTAL PER<br>STRUCTURE | 2.25 ± 0.04 | 5.76 ± 0.06 | 7.80 ± 0.03 | 3.80 ± 0.04 |

### Water dynamics around peptide bonds

To investigate the likelihood of a given peptide bond to undergo hydrolysis, the water residence times along the peptide chain were evaluated. Water molecules that linger around the peptide for too little time or for too long are unlikely to react^19^. In the first case, the contact is short-lived and there is probably not enough time for the molecule to engage in hydrolysis. In the second case, the hydrogen bonds between the water molecule and the peptide bond have a stabilising role and it is therefore not energetically favourable for the peptide bond to hydrolyse. An intermediate water residence time would then be ideal for a reaction to occur. To determine these ranges, the lifetimes for hydrogen bonds between a water molecule and the atoms in a peptide bond (C O N H) were determined, as a proxy for water residence times. For that purpose, the corresponding hydrogen bond autocorrelation functions for each peptide bond-water interaction were calculated for the ovine and the caprine β-lactoglobulin T_125_PEVDXEALEK_135_ peptides.

The decay functions are represented in Figure 4 and the average time it takes for it to reach zero is listed in Table 4. For both peptide variants, most decay functions reach zero between approximately 300 and 600 ps. These have been established as intermediate residence times^19^ and these water molecules could therefore engage in hydrolysis. A few peptide bond-water hydrogen bonds decay quite rapidly. They consist of interactions with the Thr_1_-Pro_2_ bond (Structure 3s of the ovine peptide and Structure 1g of the caprine peptide), and with the Glu_7_-Ala_8_ bond (Structure 2s of the ovine peptide). These peptide bonds are unlikely to react. As for the longer times, only two peptide bonds have decay functions that take more time to reach zero and may not react: the Glu_3_-Val_4_ and Ala_8_-Leu_9_ in Structure 4g of the caprine T_125_PVED**K**EALEK_135_ peptide. However, it is important to note that these hydrogen bond lifetimes still tend to zero at longer times. By contrast, the hydrogen bond autocorrelation functions of structural water molecules in the bovine variant of the β-lactoglobulin peptide, T_125_PVED**D**EALEK_135_, tend to a constant C(t) value^19^, which indicates the stability of these water molecules in time (see Figure S1 for an example). The water molecules around the Glu_3_-Val_4_ and Ala_8_-Leu_9_ peptide bonds in Structure 4g of T_125_PVED**K**EALEK_135_ might be then able to engage in hydrolysis, despite their autocorrelation function taking longer to reach zero when compared to the autocorrelation functions of other hydrogen bonds in the same peptide. In these cases, it is then necessary to evaluate the likelihood of these peptide bonds to undergo hydrolysis from different perspectives. These will be further discussed in the following section.

**Figure 4.**
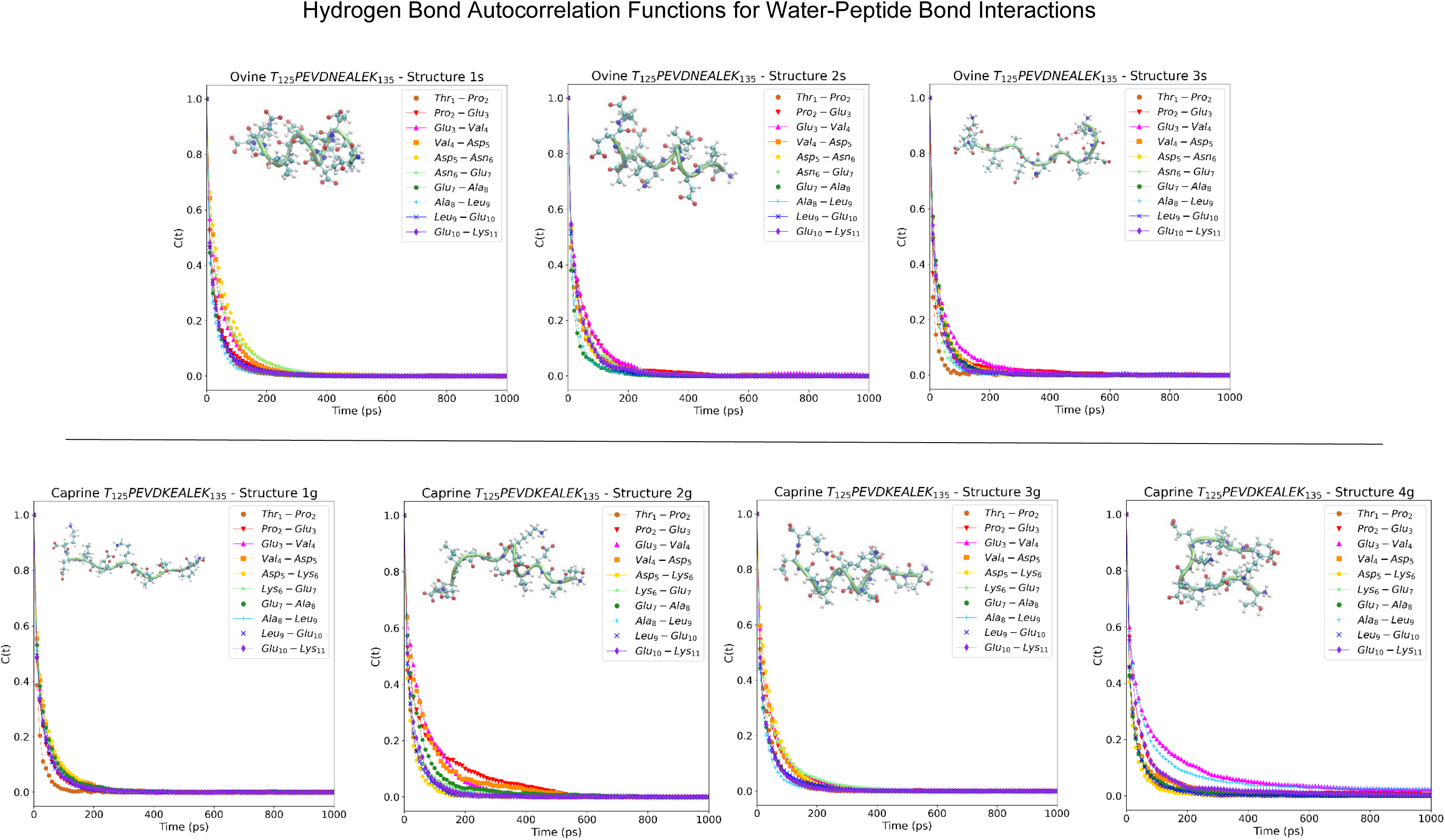
Hydrogen bond autocorrelations functions (C(t) for the peptide bond -- water interactions.

**Table 4.** Hydrogen bond autocorrelation function decay time ranges for the ovine and the caprine β-lactoglobulin peptides. An inferior and superior average time limit is showcased for intermediate decay times. Any hydrogen bond with decay times within the intermediate range might represent water molecules suitable for hydrolysis.

| Ranges | Ovine T<sub>125</sub>PVEDNEALEK<sub>135</sub> | Caprine T<sub>125</sub>PVEDKEALEK<sub>135</sub> |
| --- | --- | --- |
| Fast decay times | 220.0 ± 0.0 ps | 240.0 ± 50.0 ps |
| Intermediate decay times | 392.3.0 ± 54.4 ps 636.9 ± 129.9 ps | 420.0 ± 38.5 ps 685.0 ± 116.6 ps |
| Long decay times | -- | 2060.0 ± 267.6 ps |

As previously established, none of the stable structures found for the ovine and caprine BLG peptides rely on Na^+^ as a structural component. Therefore, the ions in solution do not influence the availability of water molecules around acidic residues in the peptide chain (Figure S2). Contrary to what is observed for some of the structures of the bovine BLG peptide in solution at neutral pH, all aspartates and glutamates are then able to participate in reaction mechanisms for peptide bond hydrolysis that involve the side chain carboxylate in all stable conformations of the ovine and the caprine BLG peptides.

### Potential sites for hydrolysis

For hydrolysis to take place, the atoms in the peptide bond (C O N H) and the surrounding water molecules need to be at an optimal geometric arrangement for the reaction to occur. Moreover, water molecules need to linger around the reaction site for long enough to react. Several authors have attempted to uncover the reaction mechanism of peptide bond hydrolysis^20–26^ and a few key geometric features have been reported^20,21,25^. Although these studies were conducted using smaller model compounds, such as formamide, they still provide a guide to identifying potential reactant structures along the amino acid chain, as previously established by our group (see Fonseca et al^19^). No geometric arrangements that matched the parameters published by Antonczak et al.^21^ and Pan et al.^25^ were detected in the ovine and in the caprine peptides (Figure S3). Therefore, the following discussions will only focus on the reactant structure published by Gorb et al^20^.

Literature values for the optimal peptide bond-water arrangement in a pre-reactive complex have a carbonyl oxygen to water hydrogen distance (C=O--H_w_) of approximately 1.8 Å and an amine hydrogen to water oxygen distance (NH--O_w_) between 1.8 Å and 2.0 Å^20^. For both the ovine T_125_PEV**N**EALEK_135_ and the caprine T_125_PEVD**K**EALEK_135_ peptides, the radial distribution functions (RDFs) for the carbonyl oxygen (C=O) and water hydrogen (H_w_) give a first maximum at approximately 1.8 Å, consistent with the reported value by Gorb et al^20^. The limiting factors to identifying potential sites for hydrolysis then become the NH--O_w_ distance and the residence time of water molecules around the carbonyl group, as it is a primary site of attack for hydrolysis to occur^20–22,24,25^. Analogous to the analysis performed for the bovine T_125_PEVD**D**EAELK_135_ peptide in solution^19^, the average number of hydrogen bonds around each carbonyl was obtained by integrating the C=O--H_w_ RDF and plotting against their respective hydrogen bond lifetimes. Figures 5A and 6A show the resulting plots, for the ovine and the caprine peptide, respectively. The plots were then split into quadrants, where the top right quadrant represents peptide bonds with a suitable amount of water molecules with adequate hydrogen bond lifetimes to react. Although the value for the horizontal threshold is straightforward as at least one water molecule will always be needed for reaction to occur, the vertical threshold depends on the distribution of hydrogen bond lifetimes for each of the peptides. For both peptides in this study, these distributions can be found in Figure S4. The average value given by intermediate hydrogen bond lifetimes is 30 ± 10 ps for the ovine peptide, and 33 ± 15 ps for the caprine peptide. In both cases, they fall within error of the reported value of 39 ± 4 ps for the bovine peptide^19^. Therefore, the vertical threshold of 39 ps was maintained for comparative purposes. Finally, Figures 5B and 6B show the average values for the NH--O_w_ distance, the second geometric parameter of the reactant structure of Gorb et al.^20^, for the ovine and the caprine peptides, respectively.

**Figure 5.**
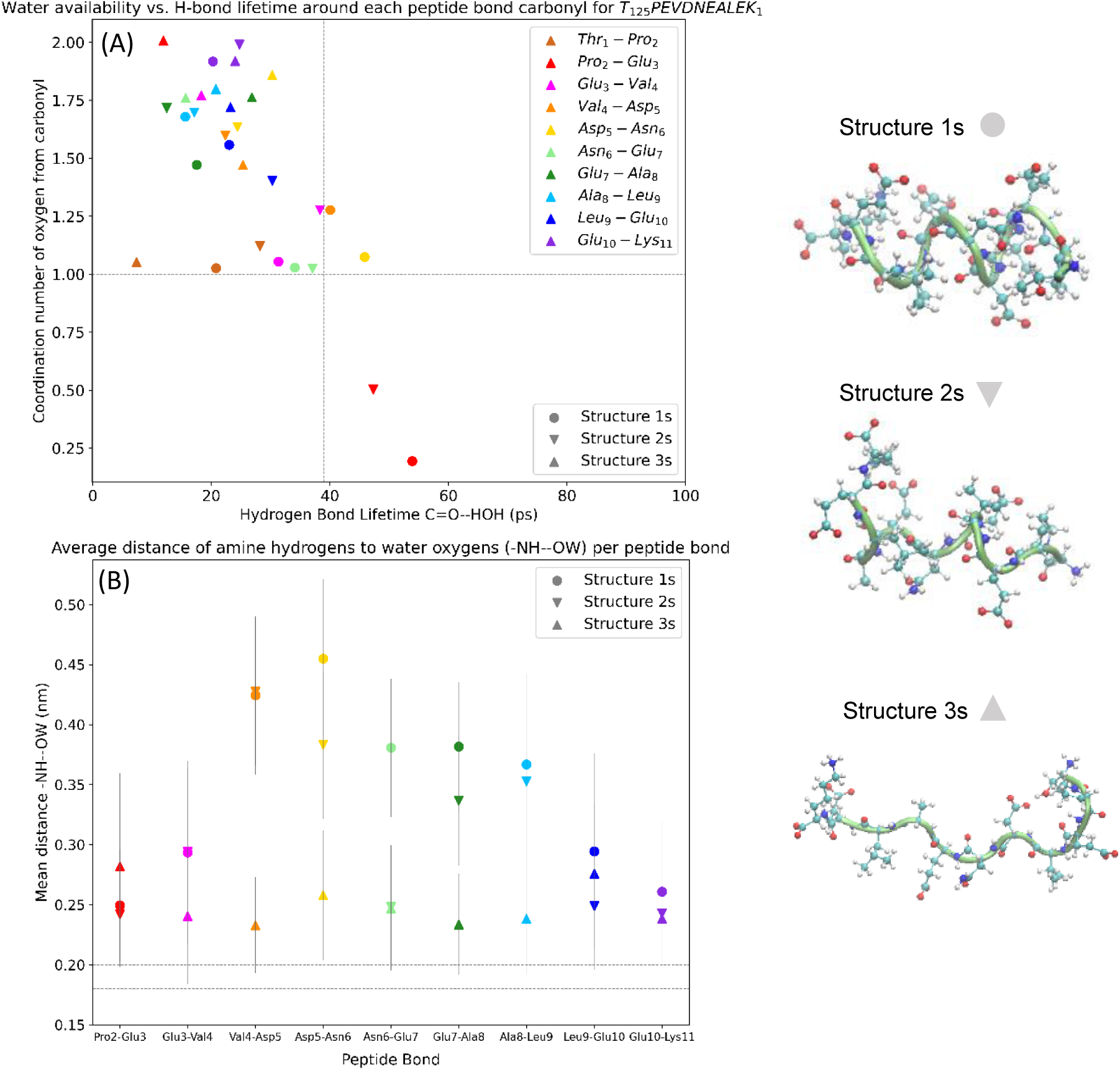
Coordination number of the carbonyl oxygen versus hydrogen bond lifetimes for C=O--H_w_ interactions (A) and average distance between amine hydrogen (NH) and water oxygen (O_w_) (B) for the ovine T_125_PEVD**N**EALEK_135_ β-lactoglobulin peptide.

**Figure 6.**
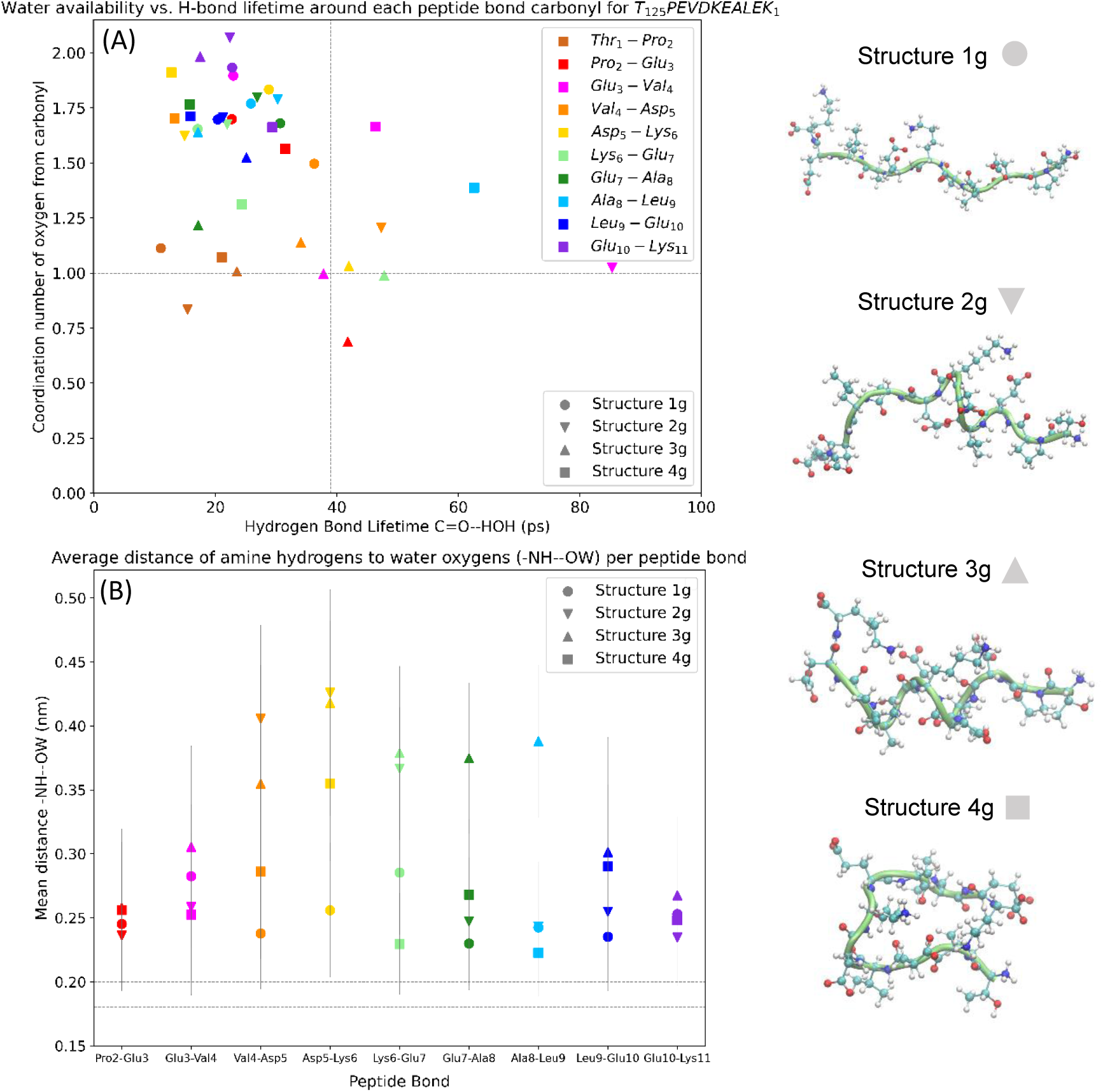
Coordination number of the carbonyl oxygen versus hydrogen bond lifetimes for C=O--H_w_ interactions (A) and average distance between amine hydrogen (NH) and water oxygen (O_w_) (B) for the caprine T_125_PEVD**K**EALEK_135_ β-lactoglobulin peptide.

In the case of the ovine T_125_PEVD**N**EALEK_135_ peptide, no peptide bonds in any of the structures have a suitable geometric arrangement with adequate water residence times to react, despite several peptide bonds having C=O--H_w_ and NH--O_w_ distances that match literature values^20^. This suggests a particular robustness of T_125_PEVD**N**EALEK_135_ in solution at neutral pH. Given that the carbonyl is the primary site of attack for hydrolysis, the water dynamics solely around this group was considered separately. In this case, only the helix formation (Structure 1s) has two peptide bonds with water molecules that could linger for long enough to undergo hydrolysis. They are the bonds Val_4_-Asp_5_-Asn_6_ located from the middle towards the N-terminal of the amino acid chain. In these cases, the carbonyl oxygen, instead of hydrogen bonding with the rest of the peptide, is exposed and vulnerable to attack.

In the caprine T_125_PEVD**K**EALEK_135_ peptide, two peptide bonds have geometric parameters and water residence times suitable for hydrolysis to occur. These are the Glu_3_-Val_4_ and Ala_8_-Leu_9_ in Structure 4g. No peptide bonds with a suitable combination of parameters and hydrogen bond lifetimes are found in any of the other structures for the caprine peptide. When considering only the water dynamics around the carbonyl group, the caprine peptide follows the same trend as the ovine peptide, with the most vulnerable bonds located from the middle towards the N-terminal side of the chain. These include the sequence Glu_3_-Val_4_-Asp_5_-Lys_6_-Glu_7_.

It is important to note the effect of structure on the vulnerability of peptide bonds in the β-lactoglobulin peptides. Table 5 summarises the exposed peptide bonds in each of the conformations for each of the β-lactoglobulin peptides. For the ovine and caprine peptides, most bonds that are likely to hydrolyse are located in helix or partial helix structures and towards the N-terminal of the amino acid chain. The only exception is the Ala_8_-Leu_9_ peptide bond in Structure 4g of the caprine peptide. Structure 4g is a loop-type structure, similar to several of the stable structures detected for the bovine peptide^19^. In that instance, the Ala_8_-Leu_9_ peptide bond was also found as a vulnerable bond for the bovine T_125_PEVD**D**EALEK_135_ in solution at neutral pH^19^. Perhaps then this is a particularly exposed bond for loop-type structures in the β-lactoglobulin peptides. Additionally, it was determined that these loop conformations are more likely to hydrolyse from the C-terminal inwards^19^. This trend is reversed here, as helix and partial helix formations seem more prone to hydrolyse towards the N-terminal instead. In the same fashion, the only vulnerable bonds towards the N-terminal that were detected in the bovine peptide were also in partially stretched conformations (similar to a partial helix), or in a stretched conformation that was stabilised by Na^+^ ions^19^. Interestingly, none of the stretched structures for the ovine and the caprine variants of the peptide display peptide bonds that are likely to hydrolyse based on the criteria determined in this study, despite having higher degrees of freedom than folded or partially folded structures. This is probably because water molecules are not held for long enough around their peptide bonds to react. Such an effect is likely due to a lack of simultaneous interaction of a given water molecule with different regions of the amino acid chain (or Na^+^ in the case of the bovine peptide), that could help in retaining them in the vicinity of a peptide bond for longer periods of time.

**Table 5.** Summary of vulnerable peptide bonds in each of the structures of the ovine and the caprine variants of the β-lactoglobulin peptide. Bonds that have adequate geometric parameters (as per Gorb et al.) and water residence times are highlighted in bold. Data for the total number of vulnerable bonds for the bovine peptide was taken from Fonseca et al^18^.

| PEPTIDE VARIANT | CONFORMATION | VULNERABLE PEPTIDE BONDS | TOTAL NUMBER OF VULNERABLE BONDS |
| --- | --- | --- | --- |
| **OVINE**<br>**T<sub>125</sub>VEDNEALEK<sub>135</sub>** | Structure 1s (helix) | Val<sub>4</sub>-Asp<sub>5</sub>-Asn<sub>6</sub> | 2 |
|  | Structure 2s (partial helix) | ----- |  |
|  | Structure 3s (stretched) | ----- |  |
| **CAPRINE**<br>**T<sub>125</sub>PEVDKEALEK<sub>135</sub>** | Structure 1g (stretched) | ----- | 6 |
|  | Structure 2g (partial helix) | Glu<sub>3</sub>-Val<sub>4</sub>-Asp<sub>5</sub> |  |
|  | Structure 3g (helix) | Asp<sub>5</sub>-Lys<sub>6</sub>-Glu<sub>7</sub> |  |
|  | Structure 4g (loop) | **Glu<sub>3</sub>-Val<sub>4</sub>, Ala<sub>8</sub>-Leu<sub>9</sub>** |  |
| **BOVINE**<br>**T<sub>125</sub>PEVDDEALEK<sub>135</sub>** | Structure 1c (folded) | Glu<sub>7</sub>-Ala<sub>8</sub>, Leu<sub>9</sub>-Glu<sub>10</sub>-Lys<sub>11</sub> | 8 |
|  | Structure 2c (folded) | Leu<sub>9</sub>-Glu<sub>10</sub> |  |
|  | Structure 3c (folded) | **Ala<sub>8</sub>-Leu<sub>9</sub>** |  |
|  | Structure 4c (stretched) | Pro<sub>2</sub>-Glu<sub>3</sub>, Asp<sub>6</sub>-Glu<sub>7</sub> |  |
|  | Structure 5c (stretched) | Val<sub>4</sub>-Asp<sub>5</sub> |  |

The mutant amino acid in position 6 of the chain has a clear effect on the types of structures the peptide can adopt, which translates in differences in the vulnerability of peptide bonds along the chain. Considering then the total amount of bonds that are likely to hydrolyse in each peptide (see Table 5), the bovine T_125_PEVD**D**EALEK_135_ β-lactoglobulin is probably more vulnerable than the other peptides in solution at neutral pH. Conversely, the ovine T_125_PEVD**N**EALEK_135_ peptide, with only two vulnerable bonds, is then the most resistant to hydrolysis in the same conditions. In whole proteins, loop regions are known to be more prone to attack then secondary structures^10,11,13^, given their high solvent exposure and their high flexibility, which enables them to adopt peptide-water arrangements that are apt to hydrolyse more easily. It is then not surprising that the bovine T_125_PEVD**D**EALEK_135_ peptide is the most vulnerable, when its behaviour is compared to that of the ovine and the caprine peptides in solution at neutral pH.

## Conclusion

The dynamics of the ovine T_125_PEVD**N**EALEK_135_ and the caprine T_125_PEVD**K**EALEK_135_ β-lactoglobulin peptides was evaluated in bulk water at neutral pH, and compared to the results previously obtained for the bovine T_125_PEVD**D**EALEK_135_ peptide. The outcome allows for the conclusion that two requirements need to be fulfilled for preservation of T_125_PEVDXEALEK_135_ to occur: reduced degrees of freedom - that impedes the peptide from adopting conformations suitable for hydrolysis -, and reduced water availability.

Peptide structure has a direct influence on whether, and how well, these requirements are fulfilled. The mutant amino acid and the charge distribution along the chain dictate the types of structures T_125_PEVDXEALEK_135_ preferentially adopts for each of the variants. The bovine peptide T_125_PEVD**D**EALEK_135_, for example, has a high concentration of negative charge in the middle of the chain, that contributes to its flexibility and ease to change between a loop-type folded structure and an unfolded structure, both of which may rely on structural Na^+^ or water molecules to stabilise the repulsive forces caused by the sequential acidic groups. By contrast, the ovine (T_125_PEVD**N**EALEK_135_) and caprine (T_125_PEVD**K**EALEK_135_) peptides have a neutral (Asn) and a positively charged residue (Lys) in the middle of the chain, respectively, which already help offset the repulsion between the side chains of aspartates and glutamates. Unlike the bovine peptide, they do not rely on interactions with Na^+^ for added stabilisation. These peptides preferentially form helices, partial helices and stretched conformations. In addition, the ovine peptide has a clear preference for a folded helix type formation, whereas the caprine peptide has a slight advantage towards unfolded structures. In general, loops and stretched conformations are more prone to hydrolysis given a higher exposure to water and more degrees of freedom, which allow them to adopt suitable arrangements to react. Helices, on the other hand, have a higher structuring and lower solvent accessibility, which makes them less vulnerable to attack. Not coincidentally then the number of vulnerable peptide bonds increases from ovine and caprine - which are able to adopt helix conformations - to caprine and bovine, which are able to adopt loop and stretched conformations. This finding suggests that, in solution at neutral pH, the bovine peptide T_125_PEVD**D**EALEK_135_ is more likely to undergo hydrolysis, followed by the caprine peptide T_125_PEVD**K**EALEK_135_ and finally the ovine peptide T_125_PEVD**N**EALEK_135_, which is probably the most robust of the three in these conditions.

Further studies on the dynamics of these β-lactoglobulin peptides on a mineral surface are necessary to further understand the differences in stability caused by the substitution in the middle of the chain, their vulnerability to hydrolysis and, consequently, their unlikely survival in the archaeological record.

## Acknowledgements

Beatriz Fonseca was funded by the Danish National Research Fund DNRF128 and the European Union’s Horizon 2020 research and innovation programme under the Marie Skłodowska-Curie grant agreement No 801199. Colin Freeman and Matthew Collins have also received support from the Danish National Research Fund DNRF128. In addition, Colin Freeman would like to thank funding from an EPSRC Programme Grant (Grant EP/R018820/1). For the purpose of open access, Matthew Collins has applied a Creative Commons Attribution (CC BY) license to any Author Accepted Manuscript version arising from this submission.

