## Supplementary Information for "The Survival of β-lactoglobulin Peptides in the Archaeological Record: Vulnerability vs. Sequence Variation"

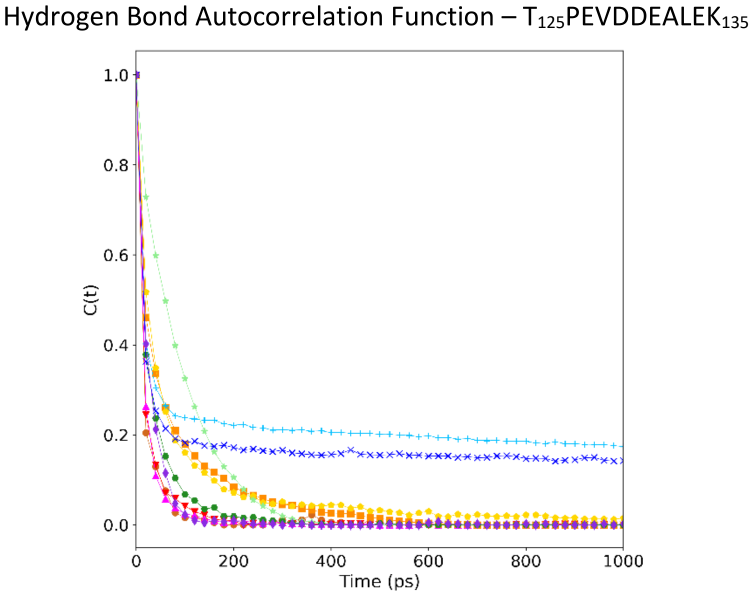


**Figure S1.** Hydrogen bond autocorrelation function for a bovine β-lactoglobulin peptide. Published in Fonseca et al[18].

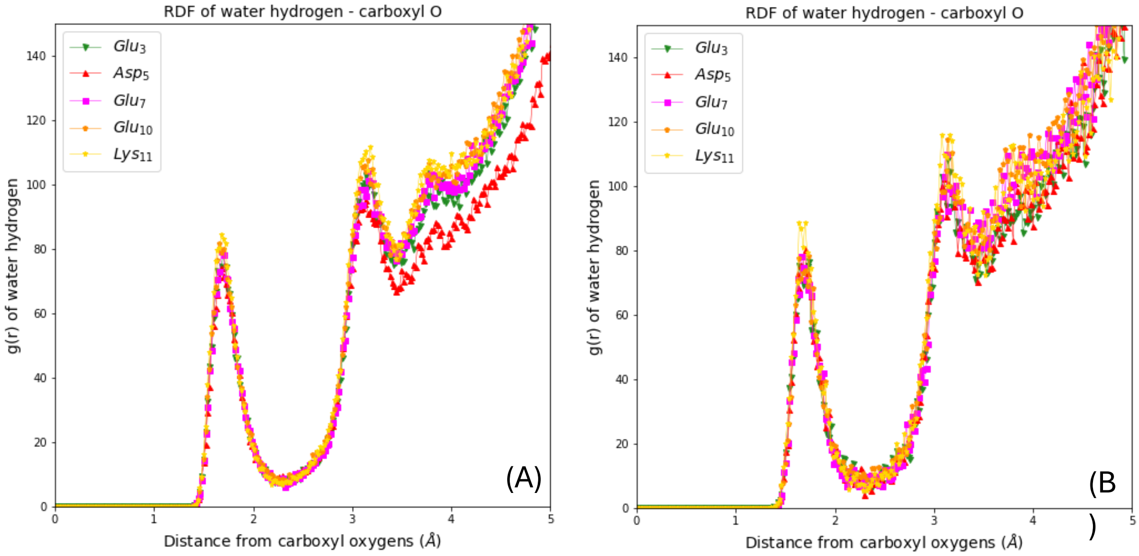


**Figure S2.** Radial distribution functions (RDF) of the carboxyl oxygen in charged residues to water hydrogen for the ovine $T_{125}PEVDNEALEK_{135}$ peptide (A) and the caprine $T_{125}PEVDKEALEK_{135}$ peptide (B). For clarity, only RDFs for Structures 1s and 1g are represented as other structures have a very similar profile.

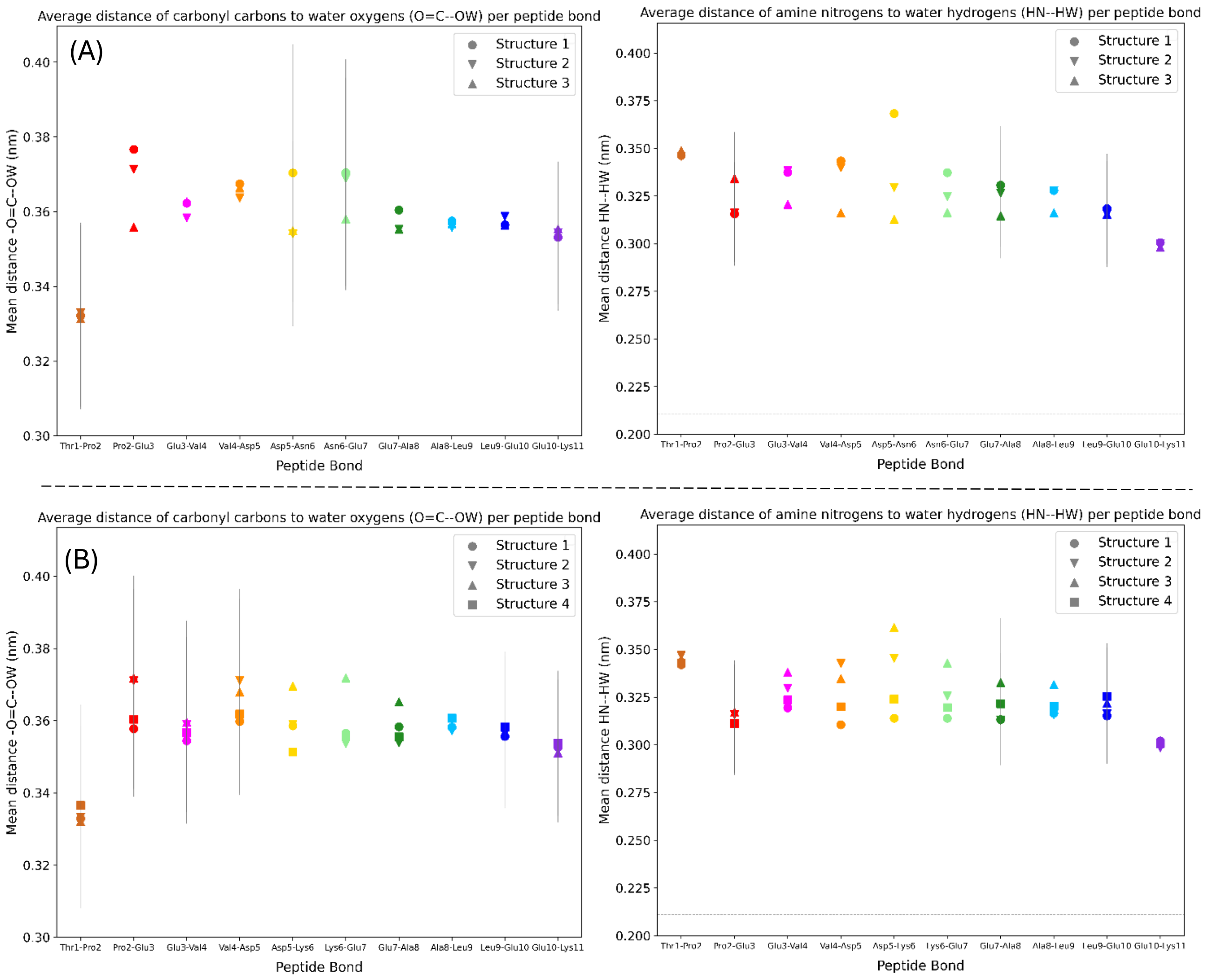


**Figure S3.** Geometric parameters for the reactant structures of Antonczak et al.[20] and Pan et al.[24] for the (A) ovine $T_{125}$PEVD**N**EALEK$_{135}$ and the (B) caprine $T_{125}$PEVD**K**EALEK$_{135}$ β-lactoglobulin peptides.

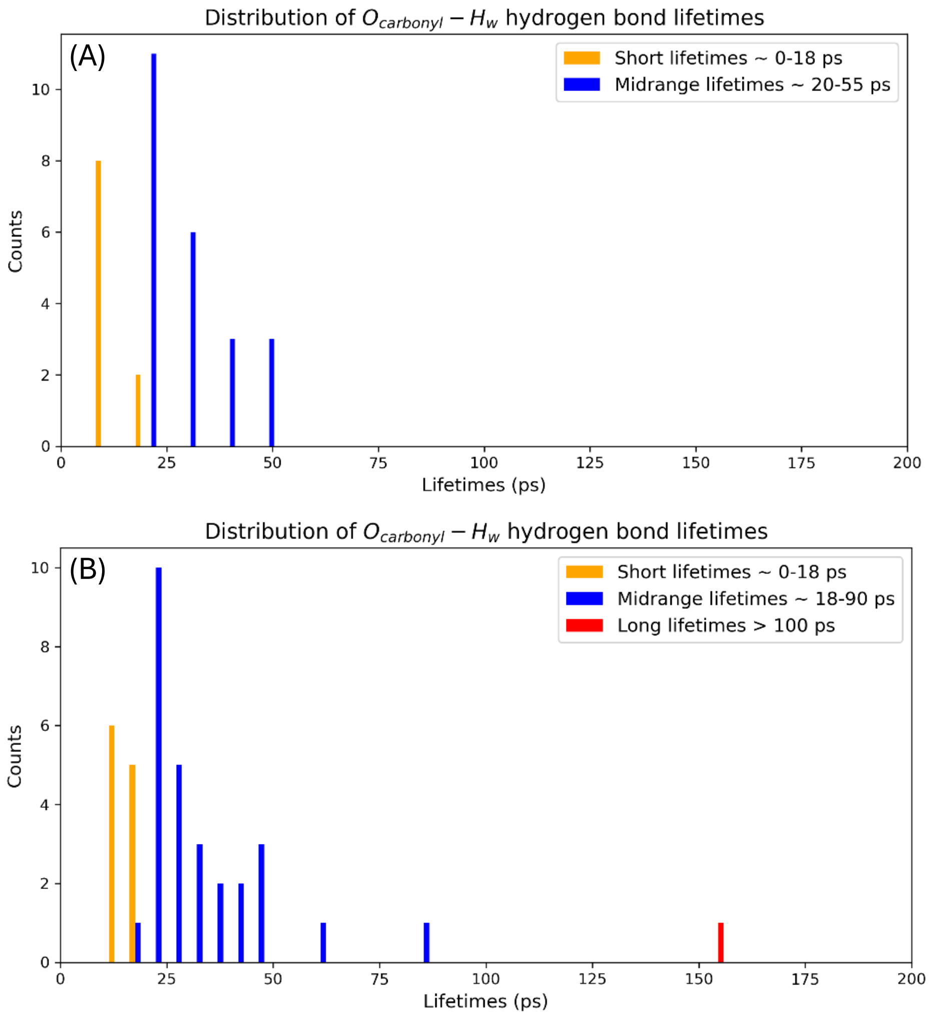


**Figure S4**. Distribution of C=O--$H_w$ hydrogen bond lifetimes for the (A) ovine $T_{125}$PEVDNEALE$K_{135}$ and (B) caprine $T_{125}$PEVDKEALE$K_{135}$ β-lactoglobulin peptides.

**Table S1.** UniProt accession numbers and PDB structure ids of the species referred to as bovine, ovine and caprine in this work.

| Peptide | UniProt Accession Number | PDB Structure id |
|---|---|---|
| **Bovine** | P02754 | 1CJ5 |
| **Ovine** | P67976 | 4CK4 |
| **Caprine** | P02756 | 4OMX |
